# Multifaceted evolution of dental morphology during the diversification of the bat superfamily Noctilionoidea

**DOI:** 10.1101/2025.06.10.658888

**Authors:** Camilo López-Aguirre, Gustavo A Ballen, Suzanne J Hand, Mary T Silcox

## Abstract

Noctilionoid bats went through one of the most extensive ecomorphological diversifications among mammals. Dietary ecology has been identified as a driver of noctilionoid morphological diversification. However, the macroevolutionary trajectories of changes dental morphology remain understudied. Studies indicate that variation in dental traits correlate with specialisation to different diets, implying differing patterns in phenotypic variability. We compared macroevolutionary trajectories across dental features quantifying five different traits using metrics of dental topography and size. Studying a sample of 110 species, we reconstructed the mode and tempo of dental evolution. We found multiple bursts of dental diversification through time, each involving different dental traits. Trait diversification was associated with different dietary radiations and could be traced to different nodes. Shifts in adaptive regimes were found in four traits, all of them concentrated within family Phyllostomidae. Evolutionary rate covariation differed across traits. We found low evolutionary covariation between measures of dental size and topography. Evolutionary modelling indicated dental traits evolved under different modes, signalling independent evolutionary trajectories. Support for diet-based models of stabilising and stochastic evolution across traits highlights the overarching effect of diet during dental evolution in Noctilionoidea. Our results support a complex and multifaceted model of evolution during noctilionoid dental morphological diversification.

## Introduction

Ecological interactions and adaptations can influence evolutionary trajectories and alter diversification processes (i.e. eco-evolutionary dynamics; Pelletier et al., 2009). As species encounter different ecological opportunities and constraints, selective forces will change, repatterning the mode and tempo of evolution (Muñoz et al., 2018). Phenotypic changes are crucial for the adaptability of species and are also informative proxies to understand underlying mechanisms shaping biodiversity (Fisher, 1930; Hallgrímsson et al., 2012; Hallgrimsson & Hall, 2005). Moreover, not only is phenotypic variation a driver of evolution, but itself is also driven by other factors that influence its magnitude and direction (Bell et al., 2011; Makedonska et al., 2012; Porto et al., 2008).

Accelerated speciation usually happens in tandem with increased morphological disparity (Arbour et al., 2019; Gavrilets & Losos, 2009; Stroud & Losos, 2016). Evolutionary trait covariation has been suggested to either constrain or promote phenotypic disparity, affecting the evolvability of traits (Felice et al., 2018; Goswami et al., 2014). Yet, the role that trait covariation (or the lack thereof) plays during taxonomic diversification needs further testing (Foote, 1997). Studying the basis of evolutionary covariation informs our understanding of macroevolutionary trajectories, elucidating what drives evolutionary lability (Felice et al., 2018). Recent studies have also provided evidence suggesting decoupled phenotypic variation at multiple scales (e.g. from single structure to complex functional unit) in traits commonly considered indivisible (Bell et al., 2011; Kilbourne & Hutchinson, 2019; López-Aguirre, Hand, Koyabu, et al., 2021). Independent processes of morphological variation could facilitate the evolutionary lability of traits, promoting ecological diversification and specialisation (Brocklehurst & Benson, 2021; Felice et al., 2018; Machado et al., 2018; Orkney et al., 2021; Slater & Friscia, 2019; Yuan et al., 2019). Elucidating the macroevolutionary trajectories of phenotypic covariation could better inform the link between phenotypic and taxonomic diversity, especially in major diversification events.

The bat superfamiliy Noctilionoidea includes all the dietary categories found in the order Chiroptera, despite representing less than a quarter of living species. As such, it is a good example of diversification driven by ecological opportunity (Arbour et al., 2019, 2021; Rojas et al., 2016). Within Noctilionoidea, the family Phyllostomidae has been the focus of multiple studies testing the eco-evolutionary dynamics of phenotypic variability and species diversification (Fleming et al., 2020; Monteiro & Nogueira, 2011; Nogueira et al., 2009; Rossoni et al., 2017, 2019; Santana et al., 2010, 2012; Santana & Dumont, 2009). Stemming from a putatively insectivorous ancestor, noctilionoids diversified throughout the southern hemisphere, the Americas and the Caribbean, reaching the highest dietary diversity and body-size range among bats (Arbour et al., 2019; Giannini et al., 2020; Gunnell et al., 2014; Hand et al., 2018; Rojas et al., 2016). Noctilionoid bats also exhibit diversification in roosting behaviours that might have facilitated their radiation (Rossoni et al., 2019; Santana, Dial, et al., 2011). Previous studies have found evidence pointing to diet as a major driver of the ecomorphological diversification in Noctilionoidea (Arbour et al., 2019; Dumont et al., 2012; Freeman, 1988; López-Aguirre et al., 2022; Monteiro & Nogueira, 2011; Rossoni et al., 2017, 2019; Santana et al., 2010, 2012). Macroevolutionary trajectories of cranial phenotypic integration in phyllostomid bats seem to have remained relatively stable across time, but with major shifts during the adoption of new adaptive regimes (Rossoni et al., 2019).

Compared with the range of studies on the cranial ecomorphological diversification of bats and its dietary basis (Arbour et al., 2021; Dumont et al., 2012; Monteiro & Nogueira, 2011; Rossoni et al., 2017), the dental evolutionary history of Chiroptera remains comparatively understudied (Esquivel et al., 2021; Freeman, 1988; Horáček & Špoutil, 2012; López-Aguirre, 2014; López-Aguirre et al., 2022; Santana, Strait, et al., 2011; Zuercher et al., 2020). Phylogenetic patterning and lack of an ecological signal have been identified in the distribution of dental anomalies (i.e. extra or missing teeth) among Chiroptera, suggesting an ancestral or developmental origin (Esquivel et al., 2021; López-Aguirre, 2014). Dietary and dental morphological differences in Neotropical bats have been found to correlate, suggesting functional demands during food processing represent selective pressures not only on cranial but also on dental morphology (Nogueira et al., 2009; Santana et al., 2010; Santana, Strait, et al., 2011). Changes in tooth size and dental proportions in the phytophagous Old World bats (Pteropodidae) have been associated with body size and phylogenetic relationships (Freeman, 1988; Zuercher et al., 2020). So far, most studies have typically focused on the size aspect of dental morphology (Self, 2015; Zuercher et al., 2020), with less emphasis on other properties of bat dental morphology (Pineda-Munoz et al., 2017).

Dental topographic analysis (DTA) has emerged as a comprehensive approach to investigate different dental morphofunctional properties across a wide range of taxa (Berthaume et al., 2020; Boyer, 2008; Evans et al., 2007; López-Aguirre, Czaplewski, et al., 2021; Peredo et al., 2018; Pineda-Munoz et al., 2017; Selig et al., 2019, 2021; Ungar & M’Kirera, 2003). Although most comprehensively applied to study Primates, both living and extinct (Allen et al., 2015; Cooke, 2011; Ledogar et al., 2013; López-Torres et al., 2018; Prufrock et al., 2016; Winchester et al., 2014), it has proven informative in studies of other mammals (Peredo et al., 2018; Pérez-Ramos et al., 2020; Pineda-Munoz et al., 2017; Selig et al., 2019) and other tetrapod groups (Melstrom, 2017; Melstrom & Irmis, 2019). Few studies have applied DTA to study bats (but see López-Aguirre, Czaplewski, et al., 2021; López-Aguirre et al., 2022; Santana, Strait, et al., 2011). Santana et al. (2011) recovered a strong ecological signal in DTA of phyllostomid bats, revealing higher dental complexity in frugivorous species. López-Aguirre et al. (2021) used DTA to compare dental morphology in modern and extinct noctilionoid bats to estimate the diet of a fossil Neotropical bat. Finally, López-Aguirre et al. (2022) found diet-based patterns of dental morphological variation, identifying multiple patterns of dental ecomorphological specialisation in Noctilionoidea. For example, tooth size was found to strongly correlate with specialisation for carnivory, whereas tooth sharpness correlated more with nectarivory, and tooth crown height and curvature correlated with frugivory (López-Aguirre et al., 2022). Omnivorous and frugivorous taxa were found to exhibit disproportionately high degrees of DTA diversity (López-Aguirre et al., 2022), reflecting the proposed role that frugivory and omnivory had in the evolution of Noctilionoidea (Davies et al., 2020; Hall et al., 2021; Potter et al., 2021; Rojas et al., 2011, 2012, 2018). The differential association between DTA metrics and dietary strategies in Noctilionoidea could be evidence that low evolutionary covariation plays a role in the adaptive radiation of this group, and provides a mechanistic framework to test competing evolutionary hypotheses.

The objective of this study is to test the independence of evolutionary trajectories across different dental traits (using DTA and molar size metrics as proxies). Given the proposed importance of diet during the adaptive radiation of Noctilionoidea (Rojas et al., 2011, 2012) and the high disparity in noctilionoid dental morphology (López-Aguirre et al., 2022), we hypothesise that the tempo and mode of evolution will vary across dental traits, reflecting independent processes. We reconstructed and compared patterns of disparity through time across dental traits, assessing whether temporal trajectories of phenotypic diversification varied among them. To test whether dental traits evolved in association with different dietary radiations and/or clades, we used diversity decomposition analysis to identify ancestors in the phylogeny from which a high proportion of trait variation originated. We also identified shifts in adaptive optima across the noctilionoid phylogeny, comparing those across dental traits. Finally, we compared the mode of evolution across dental traits, fitting six competing evolutionary models representing a variety of scenarios. These models were applied to our multivariate dental trait dataset, and to each trait independently. Differences in modes of evolution across dental traits, associated with different dietary adaptations, might be indicative of decoupled evolutionary processes (López-Aguirre et al., 2022). We estimated the covariation in evolutionary rates between dental traits, based on the best-supported multivariate model.

## Methods

### Sample acquisition

We sampled a total of 115 specimens from 110 noctilionoid species (∼44% of species diversity), representing 60 genera and all modern families (Fig. 1). All recognised dietary guilds were sampled (carnivory, frugivory, insectivory, nectarivory, omnivory, piscivory and sanguivory), representing the entire dietary diversity in the superfamily (López-Aguirre et al., 2022; Rojas et al., 2016). The species analysed represented the superfamily’s entire distribution range, extending across Africa, Oceania, the Americas and the Caribbean. To assess the role of dietary specialisation in the evolution of dental morphology in bats, species’ dietary adaptations were described using the marginality index developed by Rojas et al. (2018), placing species along a gradient (from zero to one), from specialised herbivory to specialised faunivory. We discretised the range of marginality values and classified species in four dietary guilds: specialist herbivory (0 – 0.25), generalist herbivory (0.251 – 0.5), generalist faunivory (0.51 – 0.75) and specialist faunivory (0.751 - 1). Shi & Rabosky (2015)’s bat phylogeny was pruned to fit our sample and used for all evolutionary analyses. The taxonomic arrangement was updated following (Simmons & Cirranello, 2020). Only adult specimens with no exposed dentine in occlusal view were used. Because enamel in noctilionoids is fairly thin (Dumont, 1995), such specimens are necessarily nearly or entirely unworn, implying that wear likely had little to no effect on the results. We sampled lower first molars (m1s), as they have been found to reflect dietary specialisations and the ecomorphological diversity in Chiroptera (López-Aguirre, Czaplewski, et al., 2021; López-Aguirre et al., 2022). Also, m1s represent the point with the highest bite force along the toothrow, suggesting greater functional importance during mastication (Santana et al., 2022). Focusing on a single tooth, rather than studying entire toothrows, avoids confounding patterns of dental variation (e.g. different developmental biases in species with different dental formulas) with patterns of covariation among dental traits in ecologically-dissimilar taxa (Berthaume et al., 2020). μCT scans of lower mandibles were retrieved from Shi et al. (2018) and MorphoSource, Digimorph and the American Museum of Natural History (Supplementary Table 1). Processing of raw CT data was performed using freeware 3D Slicer and Meshlab (Cignoni et al., 2008; Fedorov et al., 2012), cropping each crown at the cervix of the tooth. 3D models of individual m1s were standardised to 10,000 faces, smoothed with three iterations of the Laplacian Smoothing filter and rotated so the occlusal surface is aligned with the positive z-direction in Meshlab, following published DTA protocols for bats (López-Aguirre et al., 2022). We assessed the effect of our smoothing protocol by reprocessing a subset of specimens (12% of our sample) using only two iterations before retrieving dental morphological traits, testing for differences in variance (ANOVAs) and mean (t tests).

**Figure 1.**
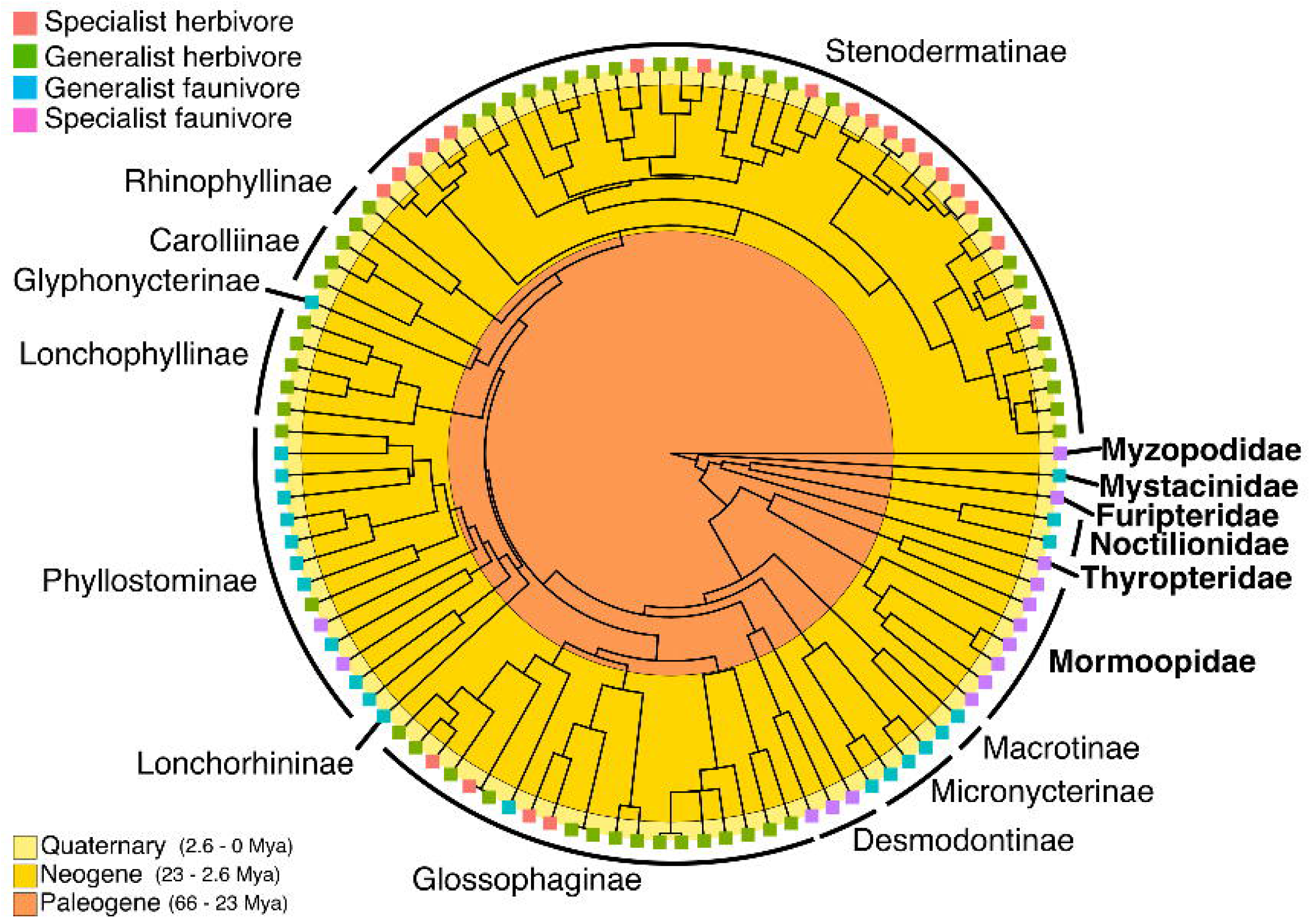
Phylogenetic hypothesis of evolutionary relationships of noctilionoid species sampled, based on a pruned version of Shi & Rabosky’s (2015) phylogeny. Tip colours indicate dietary guilds. Circular geological scale indicates times of origin.

### Dental topographic analysis

We implemented a Dental Topographic Analysis (DTA) to capture dental morphology. Unlike other commonly used morphometric methodologies (e.g. geometric morphometrics), DTA is an homology-free approach, enabling comparisons across highly divergent morphologies (Berthaume et al., 2020). Four DTA indices were calculated (Fig. 2): Dirichlet Normal Energy (DNE), Orientation Patch Count Rotated (OPCR), Relief Index (RFI), and slope. DNE quantifies the curvature of the crown to estimate its sharpness (Fig. 2A; Bunn et al., 2011), while OPCR calculates crown topographic complexity as a proxy for estimating the set of tools a tooth has to process food (Fig. 2B;Evans et al., 2007). RFI is a ratio of the crown’s 3D-to-2D surface area that acts as an index of crown height (Fig. 2C; Boyer, 2008). Slope was estimated as the mean angle of inclination across the occlusal surface of the crown (Fig. 2D), averaging the slope of all faces in the mesh (Ungar & M’Kirera, 2003). 2D area of the tooth crown (obtained to compute RFI; see Fig. 2C) was used as a measure of molar area (herein MA) (López-Aguirre et al., 2022). These five metrics have been found to be informative to explore the correlation between dental phenotypes and dietary adaptations (Berthaume et al., 2020). The metric MA only accounts for changes in size, whereas all other DTA metrics also account for form. All five metrics were calculated in the R package *molaR* (Pampush et al., 2016). We excluded the edge and top 0.1% of per-face DNE values, used an alpha value of 0.01 for estimating RFI, and set the number of steps for the OPCR rotation to 8 (step size of 5.625 degrees). DTA and MA values were log-transformed prior to all statistical analyses, standardising all metrics to allow comparison of statistical results.

**Figure 2.**
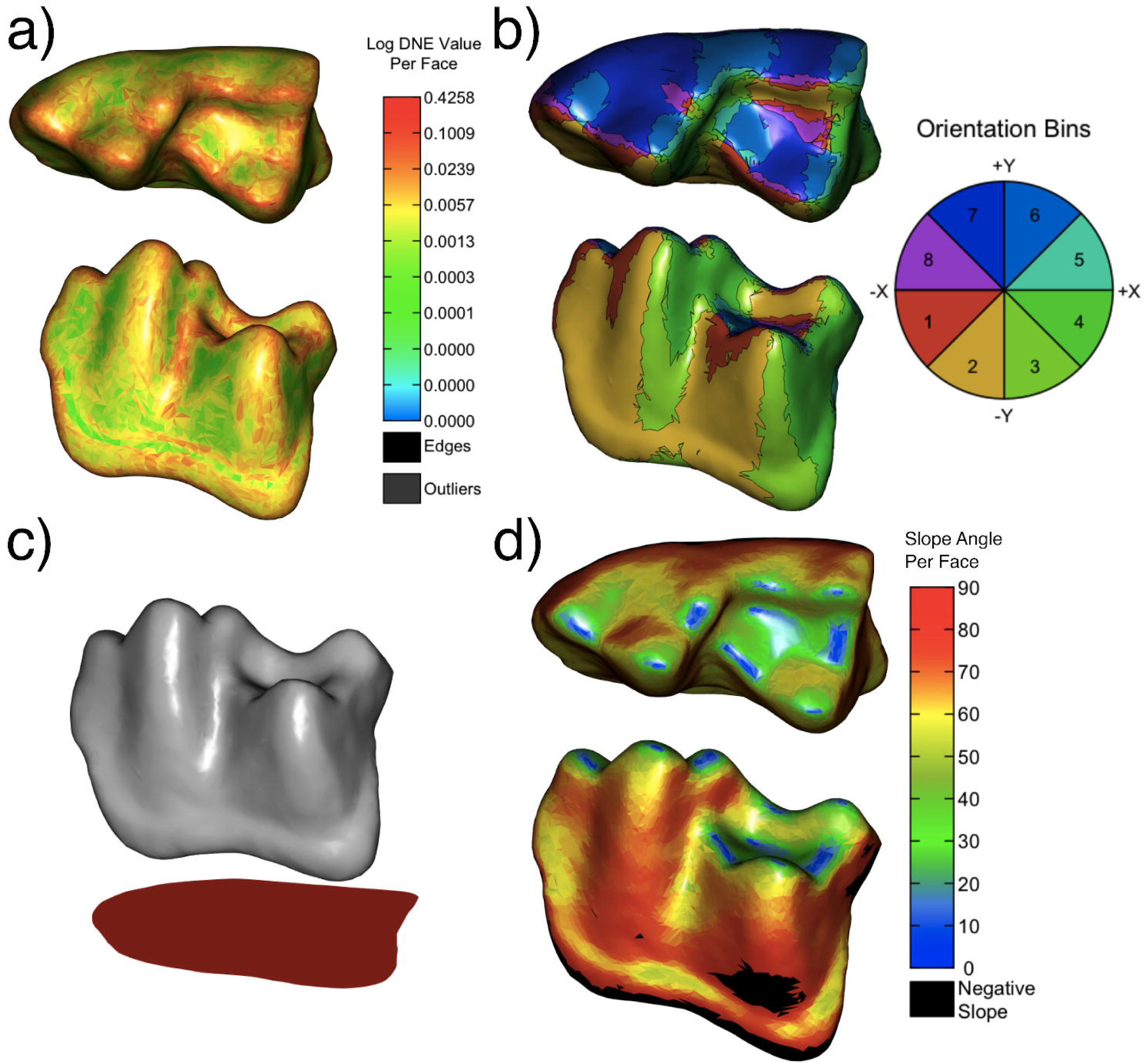
DTA metrics used to quantify four different dental traits sampled in this study, exemplified by m1 of *Micronycteris megalotis*; DNE to quantify sharpness (A), OPCR to quantify surface complexity (B), RFI to measure crown height (C), and slope (D). Molar area was quantified as the 2D area extracted for RFI (shadow in C). DNE and slope values increase from blue to red colours. Colour groupings in (B) indicate orientation of patches with respect to the 8 cardinal directions for 3D-OPCR.

We used Gol’din et al.’s (2018) protocol to codify dental wear in bats and tested for differences in variance between dental wear categories, finding no statistical support for an effect of wear on our variables (DNE: *P*= 0.061; OPCR: *P*= 0.105; RFI: *P*= 0.528; Slope: *P*= 0.949; MA: *P*= 0.282). We used phylogenetic analyses of variance (pANOVA) to test for differences in dental morphological traits between dietary guilds. Correlation patterns across these five dental traits have been reported to greatly diverge across animal groups (e.g. RFI-DNE correlation with *R^2^*= 0.76 and 0.31 in prosimians and humans, respectively; [Berthaume et al., 2020; Pampush et al., 2018; Winchester et al., 2014]). This variation implies that correlations are not simply a product of underlying mathematical similarities but are also a reflection of the biological signal in the sample.

### Reconstruction of macroevolutionary trajectories

We constructed the dental morphospace to visualise morphological differences across dietary guilds using a principal component analysis and projected our pruned phylogeny on the biplot to visualise phylogenetic relationships across species (Fig. 3). Probability kernel density functions of the four diet categories were added to our morphospace to better elucidate the partitioning of morphospace among dietary guilds. We estimated the temporal dynamics of variability in dental morphology performing disparity through time (DTT) analyses on all DTA traits, as implemented in the “dtt” function in the R package *geiger* (Pennell et al., 2014). We employed a global envelope test to calculate confidence intervals of graphical DTT while controlling for multiple tests, thereby reducing false-positive rates (Murrell, 2018). DTT compares observed trait disparity through time to that simulated under Brownian Motion (random walk process), estimating the relative subclade disparity at each internal node of the tree using the morphological disparity index (MDI; Graham J. Slater et al., 2010). Significant negative MDI values signal adaptive radiations (i.e. early burst of variation), where disparity would be highly partitioned among early-diverging clades, with more recent clades explaining smaller portions of total disparity (Slater et al., 2010).

**Figure 3.**
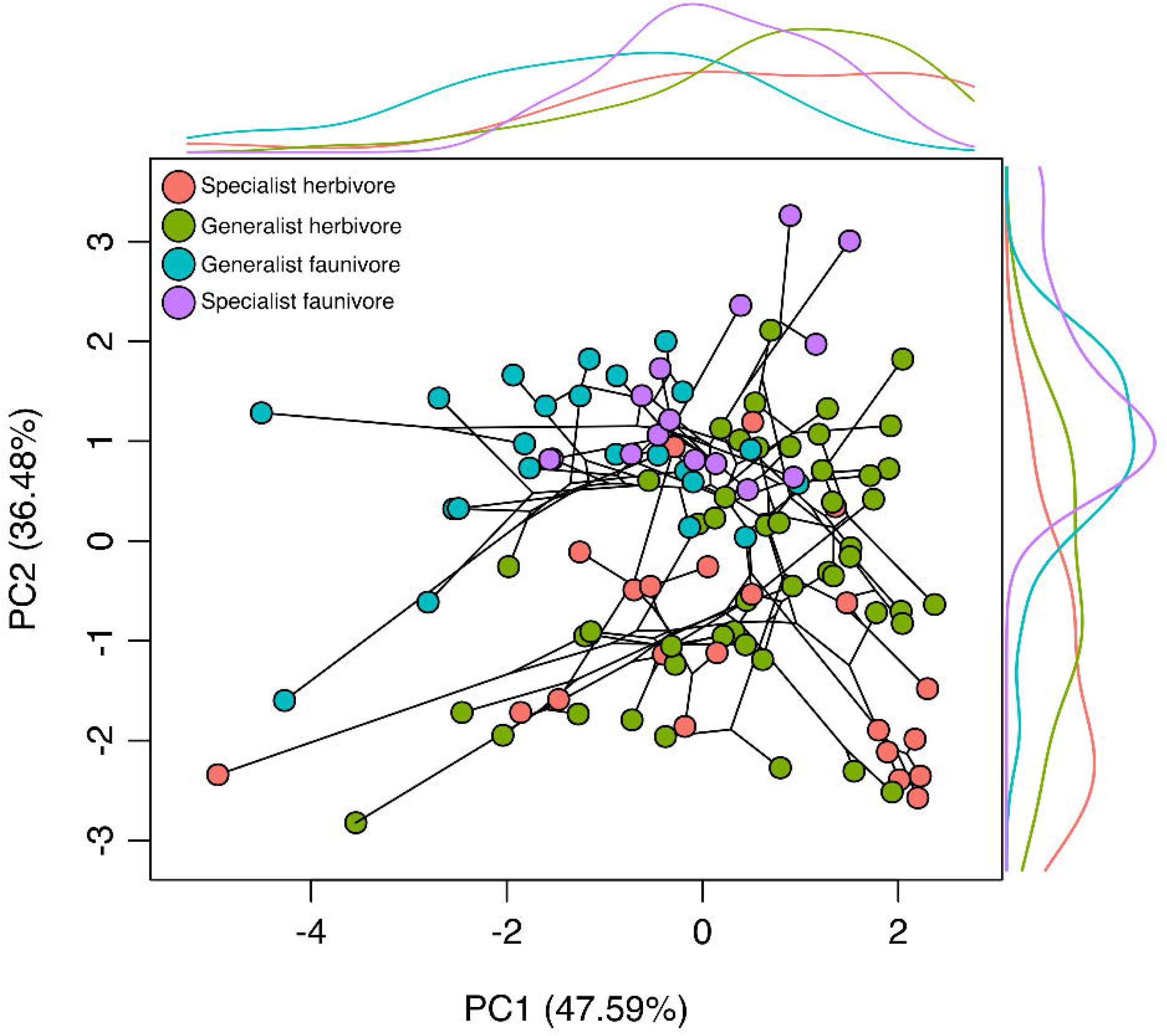
Principal component analysis reconstructing dental morphospace. Tip colours represent dietary guilds. A pruned version of Shi and Rabosky’s (2015) phylogeny was projected into morphospace. Probability density functions were plotted for each principal component, highlighting dietary structuring of morphospace.

### Dental trait diversification

We assessed whether the origin of phenotypic diversity of each dental trait is linked to the evolution of different dietary adaptations and/or clades by using a diversity decomposition analysis (DDA; Pavoine et al., 2010), as implemented in the “decdiv” function in the R package *adiv* (Pavoine, 2020). DDA identifies ancestors (nodes) in the phylogeny from which a disproportionate amount of trait diversity originates, relative to the number of descendants (tips), and provides a series of statistical tests to identify and interpret patterns found across a phylogeny (Pavoine et al., 2010). The first test (S_1_) evaluates whether a particular node accounts for the origin of most trait diversity found at the tips; the second test (S_2_) evaluates whether a few nodes account for the origin of most trait diversity, while the rest of the nodes account for little or no trait diversity (Pavoine et al., 2010). Finally, the third test (S_3_) assesses whether trait diversity originated in nodes near either the roots or the tips of the phylogeny. All three tests have a null hypothesis that nodes that gave rise to trait diversity are randomly distributed across the phylogeny (Pavoine et al., 2010). The graphical output of DDA was also exported to examine detailed patterns.

In contrast to most statistical models of trait evolution that rely on *a priori* assumptions (e.g. traits evolve similarly across an entire phylogeny), recent methodological developments allow testing of evolutionary scenarios where trait variance freely shifts across different adaptive optima (i.e. data-driven models; Uyeda et al., 2018). Data-driven approaches allow the detection of clade-specific evolutionary processes that otherwise would be obscured by over-arching evolutionary hypotheses (Uyeda et al., 2018). We identified nodes where shifts in trait optima occurred (set to a maximum of 10) in each dental trait separately, using a model of trait evolution under stabilising selection using the R package *PhylogeneticEM* (Bastide et al., 2018). Graphical outputs were used to compare patterns of adaptive shifts across the five dental traits.

### Evolutionary model testing

Previous studies have provided conflicting evidence of patterns of correlation across dental morphological traits used in this study, limiting our capacity to confidently predict how our metrics correlate (Berthaume et al., 2020; Thiery et al., 2019). To investigate this, we fitted seven competing evolutionary models: an early burst model (EB), a single-rate Brownian motion model (BM), a diet-informed multiple rate Brownian motion model (BMM), a single-optimum Ornstein-Uhlenbeck model (OU1), a diet-informed multi-optimum OU model (OU_diet_), and a version of the BM and BMM models that assumes group-specific ancestral states (BM_diet_ and BMM_diet_, respectively; O’Meara et al., 2006). Early burst (EB) models assume decreasing trait variance across time after an early rapid diversification driven by ecological opportunity, consistent with an adaptive radiation (Harmon et al., 2010). BM models assume trait variance increasing linearly across time under a single rate, proportional to lineage accumulation (Felsenstein, 1973). BMM models assume a similar scenario to a BM model, but allow for trait variance evolving under different rates on each dietary group (O’Meara et al., 2006). OU models incorporate stabilising selection, constraining trait variance towards adaptive optima across time (Hansen, 1997). OU1 assumes traits evolve in a manner that is constrained towards a single optimum and OU_diet_ assumes trait variance with dietary regimes having different adaptive optima. To fit our diet-based models (BMM, BM_diet_, BMM_diet_, OU_diet_), we first simulated the evolution of dietary guilds using stochastic character mapping, using the “make.simmap” function in the R package *phytools* (Revell, 2012). This produces a phylogenetic tree with character states (i.e. dietary guilds) and reconstructed ancestral states mapped into the phylogeny (SIMMAP tree; Bollback, 2006). These seven models were run for all five dental traits together (multivariate model) and for each trait separately. The best-fitted multivariate model enabled us to explore patterns of rate covariation across traits by estimating the stationary variance-covariance matrix using the function “stationary” in the R package *mvMORPH* (Clavel et al., 2015). The stationary covariance matrix was then standardised as a correlation matrix (removing scaling effects) and used to interpret covariance in evolutionary rates across dental traits. All models were fitted using functions “mvEB”, “mvBM” (model=“BMM” for BMM; model=“BM1” for BM) and “mvOU” (model=“OUM” for BMM; model=“BM1” for BM) in the R package *mvMORPH* (Clavel et al., 2015), and support for each model was estimated using AICc. Models with the lowest ΔAICc were considered to be supported by our data.

## Results

We found differences in temporal trajectories and modes of evolution across dental traits. The diversification of each trait was associated with the evolution of different clades and dietary adaptations. The morphospace constructed with the principal component analysis of all five dental traits revealed species clustered in two main groups, one predominantly consisting of herbivorous taxa and the other predominantly faunivores (Fig. 3). Species distribution in morphospace also revealed partitioning across the level of specialisation to either herbivory or faunivory. Generalist herbivores and faunivores separated across PC1 (explaining 47.59% of variation), whereas specialist herbivores and faunivores grouped at the opposite ends of PC2 (explaining 36.48% of variation). Phylogenetic ANOVAs revealed statistically significant differences in dental traits across dietary guilds, with the exception of slope (Table 1). T-tests and ANOVAs showed no effect of our mesh smoothing protocol in our dataset. (Supplementary Table 2).

**Table 1.** Phylogenetic ANOVAs testing for statistical differences in dental traits across dietary guilds.

|  | df | F | *P* |
| --- | --- | --- | --- |
| Sharpness | 3 | 3.7255 | 0.0169 |
| Crown height | 3 | 11.258 | <0.0001 |
| Surface complexity | 3 | 4.4343 | 0.0092 |
| Slope | 3 | 2.0473 | 0.1144 |
| Molar area | 3 | 2.8866 | 0.0428 |

Disparity-through-time analyses indicate an initial stage of evolutionary stasis during the early history of noctilionoid evolution, followed by trait-specific trajectories of diversification over time (Fig. 4). Average subclade disparity in molar area and sharpness peaked early in noctilionoid evolution, followed by a steady decrease up to the present. Occlusal surface complexity peaked at the end of the Neogene, whereas crown height and slope showed slight increases in diversity during the Quaternary. MDI values of surface complexity, crown height and slope all were significantly different from expectations under a BM model, whereas MDI values of molar area and sharpness were not (Table 2). Crown sharpness, height and slope had negative MDI values.

**Figure 4.**
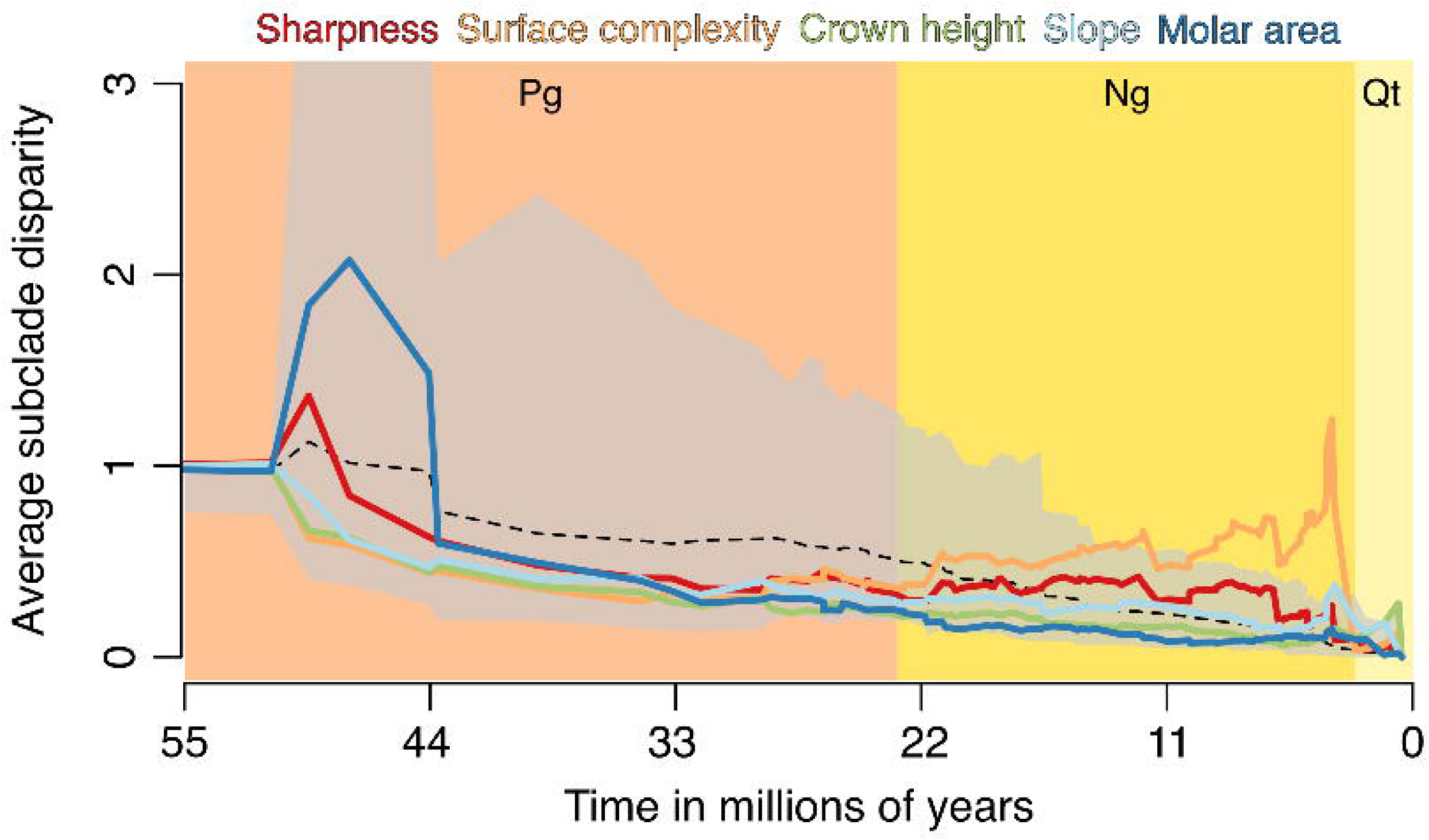
Disparity through time (DTT) plots showing temporal trajectories of dental morphological diversity. Relative time values range from zero (root of the tree) to one (present). Solid lines show the relative average disparity of subclades compared with the total clade. Dashed line shows expectation for all metrics under a Brownian motion (BM) model of evolution based on 1000 simulations. 95% confidence intervals are represented by the grey area. Sharpness is quantified with DNE, surface complexity with OPCR, crown height with RFI, crown curvature with slope and molar area with MA. Geologic periods are represented in the time axis: Paleogene (Pg), Neogene (Ng), and Quaternary (Qt).

**Table 2.**
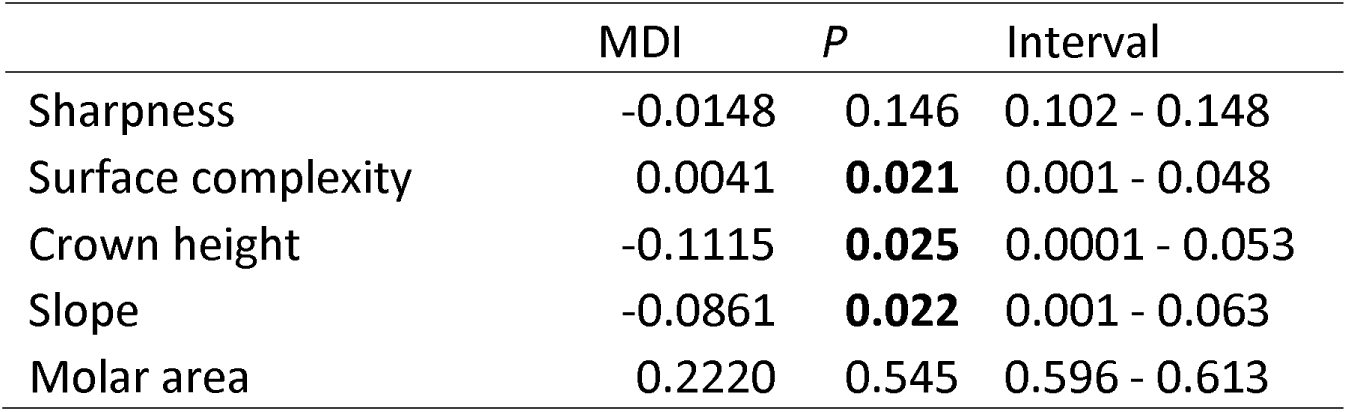
Statistical tests of disparity through time analyses for five dental traits. Morphological disparity index (MDI) compared observed values against a null hypothesis of evolution under Brownian motion (BM). *P* values and intervals are based on 1000 iterations, following the global envelope test developed by Murrell (2018). Values significant at an α=0.05 level are in bold.

Plots of diversity decomposition showed only a few ancestors at family nodes were points of origin of trait diversity (Fig. 5), and none were evident in the most basal families (i.e. Mystacinidae and Myzopodidae). A disproportionate amount of sharpness and molar area diversity originated in the most recent common ancestor of Noctilionidae, Furipteridae and Thyropteridae. Most of the diversity found across all five traits can be traced to nodes within Phyllostomidae, although each variable showed a unique pattern of change within that group (Fig. 5). Variation in sharpness can be traced to multiple nodes across all phyllostomid subfamilies, whereas no variation in surface complexity can be traced to nodes within Glossophaginae. Across all five traits, only a portion of the variation in surface complexity was disproportionally represented by the common ancestor of Desmodontinae. A high proportion of diversity in crown height can be traced to the common ancestor of all herbivorous phyllostomid subfamilies (excluding Glossophaginae). Almost no variation in slope was traced to nodes within Phyllostominae, Lonchophyllinae and Carollinae. Nodes within the faunivorous Phyllostominae subfamily and the most recent common ancestor of Noctilionidae and Furipteridae concentrate the highest proportion of diversity in molar area. Permutation tests of DDA revealed differences in patterns of statistical significance across dental traits (Table 3). Tests of statistical significance rejected the null hypothesis of the S_3_ test for crown sharpness, S_2_ and S_3_ for crown height and slope, S_1_ and S_3_ for molar area, and no null hypothesis was rejected for surface complexity.

**Figure 5.**
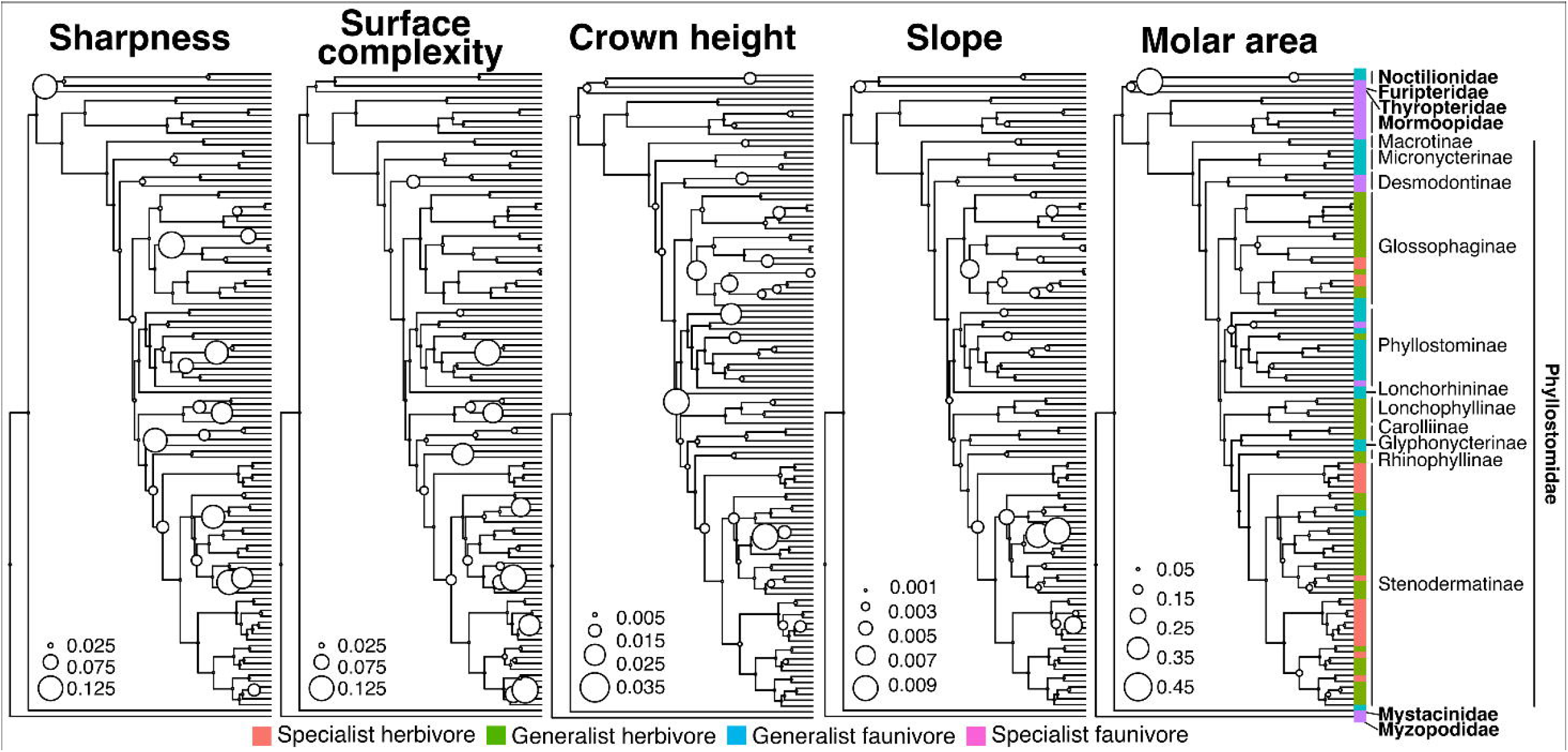
Diversity decomposition plots for five metrics showing distribution of phenotypic diversity in five dental traits across the phylogeny. Sharpness is quantified with DNE, surface complexity with OPCR, crown height with RFI, crown curvature with slope and molar area with MA. Circles represent magnitudes of dental diversity concentrated in a node and numbers indicate relative contribution to total trait diversity. Species are classified based on four dietary categories: Specialist herbivore (S herbivore), generalist herbivore (G herbivore), generalist faunivore (G faunivore) and specialist faunivore (S faunivore).

**Table 3.** Statistical tests of diversity decomposition of dental morphology for five dental traits. S_1_ tests for concentration of diversity in a single node, S_2_ tests for concentration of diversity in few nodes and S_3_ tests for skewness in concentration of diversity towards the root or tips. Values significant at an α=0.05 level are in bold.

|  | Test | Obs | Std. Obs | Alter | *P* |
| --- | --- | --- | --- | --- | --- |
| Sharpness | S1 | 0.061 | -1.2741 | greater | 0.944 |
|  | S2 | 0.545 | -0.1715 | two-sided | 0.877 |
|  | **S3** | 0.473 | 5.70214 | two-sided | **0.001** |
| Surface complexity | S1 | 0.068 | -1.3525 | greater | 0.988 |
|  | S2 | 0.544 | -1.3635 | two-sided | 0.176 |
|  | S3 | 0.373 | 1.41393 | two-sided | 0.143 |
| Crown height | S1 | 0.061 | -1.1844 | greater | 0.918 |
|  | **S2** | 0.475 | -2.4237 | two-sided | **0.012** |
|  | **S3** | 0.501 | 7.12898 | two-sided | **0.001** |
| Slope | S1 | 0.074 | -1.5259 | greater | 0.948 |
|  | **S2** | 0.49 | -3.4482 | two-sided | **0.001** |
|  | **S3** | 0.469 | 5.34342 | two-sided | **0.001** |
| Molar area | **S1** | 0.173 | 3.22904 | greater | **0.001** |
|  | S2 | 0.581 | 0.72655 | two-sided | 0.452 |
|  | **S3** | 0.517 | 6.61721 | two-sided | **0.001** |

Data-driven evolutionary modelling retrieved shifts in adaptive regimes in surface complexity, crown height, slope and molar area, with each trait experiencing regime shifts in different clades (Supplementary Fig. 1). Regime shifts were detected in the common ancestors of subfamilies Phyllostominae and Desmodontinae for surface complexity, and the common ancestors of subfamilies Glossophaginae and Stenodermatinae for crown height. Multiple regime shifts in slope were detected within Glossophaginae and Stenodermatinae, whereas a single shift in molar area was found in the common ancestor of the carnivorous bats *Vampryum spectrum* and *Chrotopterus auritus*.

Multivariate evolutionary modelling revealed OU_diet_ was the best-fitted model for our set of dental traits (Table 4). Evolutionary rate covariances derived from this multivariate OU_diet_ model revealed a strong evolutionary correlation between crown height and slope (*R^2^*= 0.85), and sharpness and surface complexity (*R^2^*= 0.79); moderately strong between molar, sharpness and surface complexity (*R^2^*= 0.46-0.57); and weak correlations (<0.3) between all other combinations of traits (Supplementary Fig. 2). Trait-specific univariate evolutionary modelling showed dental traits followed different evolutionary trajectories (Table 4). OU_diet_ was unambiguously the best-fitted model for surface complexity, slope and crown height, while it was the second-best fitted model (ΔAICc= 1) for sharpness, after BMM_diet_. BM_diet_ was the best-supported model for molar area.

**Table 4.**
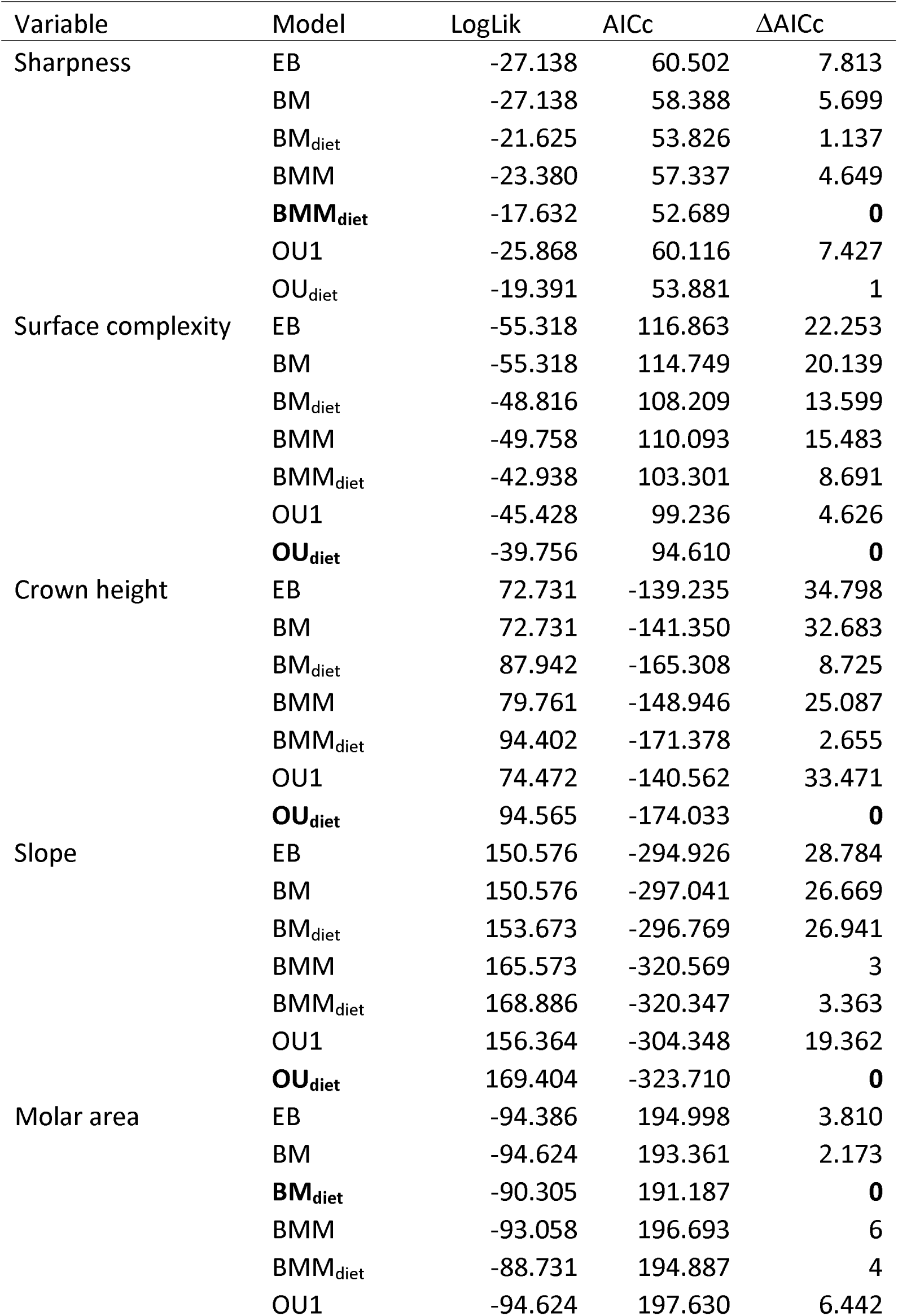
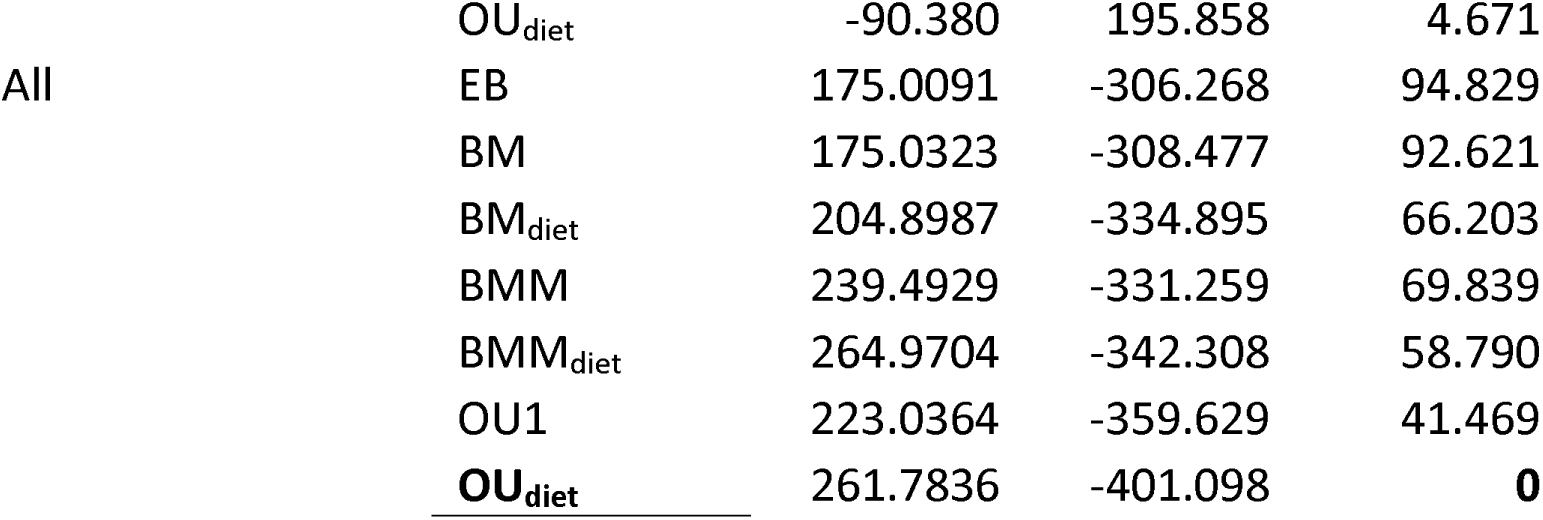
Test of competing evolutionary models for five dental traits metrics. Small sample size-corrected Akaike information criteria (AICc) was used to select the best supported models, with models with ΔAICc below two considered as supported (in bold). Univariate models were run for each trait individually, and a multivariate models was run for all traits combined.

## Discussion

Exploring the role trait covariation has shaping evolutionary trajectories provides novel insights into the mechanisms shaping diversification processes (Machado et al., 2018; Rossoni et al., 2019). Our study found that during the adaptive radiation of Noctilionoidea, different dental morphological traits showed different temporal patterns and modes of evolution, signalling independent trajectories. Our results are consistent with macroevolutionary trajectories showing decreases in cranial trait covariation during the adoption of new adaptive regimes by noctilionoids (Rossoni et al., 2019). Our results indicate a strong decoupling in macroevolutionary trajectories between dental shape (DTA metrics) and dental size (MA), and varying degrees of evolutionary covariation across dental shape traits, partially supporting our hypothesis. Disparity-through-time analyses revealed trait-specific temporal trajectories of morphological diversification, supporting our expectations. Crown height and slope showed patterns consistent with an early burst of diversification (adaptive radiation), surface complexity followed a pattern of late diversification, whereas dental size and sharpness did not deviate from a random-walk process. Dental morphological variation patterns reflected dietary specialisations and clade-specific processes across Noctilionoidea, by tracing the origin of most of the phenotypic diversity and shifts in adaptive regimes to herbivorous ancestors within Phyllostomidae (Fig. 5 and Supplementary Fig. 1). Across dental shape traits, strong covariation was found between dental sharpness (DNE) and topographic complexity (OPCR) and between crown height (RFI) and slope, but not between any other combination of traits. Evolutionary modelling signalled an overarching effect of diet during the dental morphological diversification in Noctilionoidea, with differences in processes of stabilising selection and random-walk processes across dental traits. Overall, different temporal trajectories, ancestors of origin of morphological diversification, and modes of evolution across dental morphological traits suggests labile covariation in dental traits might have played a role during the diversification of Noctilionoidea.

### Temporal trajectories of dental evolution

Temporal trajectories of dental disparity revealed multiple bursts of dental diversification across noctilionoid evolutionary history, each reflecting the diversification of different dental morphological traits. The early evolutionary stasis in trait diversity probably reflects low taxonomic and ecological diversity in Noctilionoidea, represented by small insectivorous taxa (e.g. Myzopodidae) that show little significant divergence from the ancestral state (Gunnell et al., 2014; Hand et al., 2018; Moyers Arévalo et al., 2018). A burst in molar area and sharpness disparity occurred early in noctilionoid evolution, possibly associated with the first appearance of modern families with novel dietary adaptations, like the large-bodied piscivore family Noctilionidae (Amador et al., 2018). Disparity in surface complexity and sharpness steadily increased across the Neogene, surface complexity undergoing a sharp burst in diversity by the end of the Neogene, coinciding with the adaptive radiation in Phyllostomidae and in particular with the evolution of phytophagous groups (e.g. Glossophaginae, Lonchophyllinae and Stenodermatinae; Amador et al., 2018; Fleming et al., 2020; Rojas et al., 2016). Statistical tests of DTT support our expectation of decoupled temporal trajectories of trait diversification. The random-walk process identified for molar area seems to conflict with previous studies that have highlighted the importance of shifts of increasing body size during the evolution of Noctilionoidea (Giannini et al., 2020; Moyers Arévalo et al., 2018). Reconstruction of body size ancestral states in Noctilionoidea suggests the ancestral body mass for the superfamily was 9-12 g, followed by a rapid accumulation of evolutionary shifts towards both larger (especially in carnivorous species) and smaller body sizes (Giannini et al., 2020; Moyers Arévalo et al., 2018). Considering previous studies reporting a correlation between body size and molar area in bats (López-Aguirre et al., 2022; Zuercher et al., 2020), our results suggest that shifts in the evolution of body size might have relaxed evolutionary selective forces on tooth size. Future studies should further explore the relative effect of allometry and dietary adaptations in explaining tooth size variation. Significantly negative MDI values of crown height and slope suggest an adaptive radiation of these traits driven by disruptive selection favoured by novel ecological adaptations in early diverging families, like piscivory in Noctilionidae and ground omnivorous foraging in Mystacinidae (Brooke, 1994; Hand et al., 2009).

### Trait diversity across the phylogeny

Diversity decomposition analyses revealed variation in each dental trait can be traced back to a different set of ancestors across the phylogeny, supporting our prediction of trait-specific patterns of evolution. Our results suggest high evolutionary lability in dental morphology where trait variation correlates with specific clades and dietary radiations across the phylogeny (Pavoine et al., 2010). Statistical tests of diversity decomposition revealed trait-specific patterns of significance that indicate different evolutionary trajectories of trait variation. Our results indicate that the origin of crown sharpness diversity can be traced to nodesskewed towards the root of the phylogeny. Variation in sharpness was traced to the origin of Noctilionidae and possibly the transition to piscivory in the family. The majority of increases in sharpness diversity occurred in phytophagous nodes, similar to previous diversity decomposition analyses in Phyllostomidae (Rossoni et al., 2019). Interestingly, a disproportionate amount of sharpness diversity was traced to a transition to frugivory (i.e. genus *Brachyphylla*) within the nectarivorous subfamily Glossophaginae. Higher dental sharpness in the genus *Brachyphylla* compared to its nectarivorous sister taxa has been reported (López-Aguirre et al., 2022). A nectarivory-frugivory transition could have been facilitated by increased dental sharpness from a simplified nectivore to a derived frugivore dentition.

For surface complexity, no statistical significance in any of the DDA tests indicates that it is highly labile and can evolve independently across different lineages irrespective of phylogenetic relatedness. A weak phylogenetic signal in surface complexity (i.e. OPCR) has been previously reported in Noctilionoidea (López-Aguirre et al., 2022), supporting our results indicating evolutionary lability. Most of the diversity in surface complexity can be traced to ancestors within phyllostomid, suggesting its evolutionary lability was consistent with the adaptive radiation of Phyllostomidae. The origin of crown height and slope diversity can be traced to multiple ancestors, mostly near the root of families Phyllostomidae, Noctilionidae, Furipteridae and Thyropteridae, reflecting the overall importance of these two traits in dietary adaptations and the ecological diversification of Noctilionoidea (López-Aguirre et al., 2022). Dental variations in slope and crown height have been associated with the specialisation of noctilionoids to a variety of diets, signalling they could represent labile traits for ecological evolution (López-Aguirre et al., 2022). Finally, most diversity in molar area can be traced to two ancestors involved in transitions to predation on vertebrates: the ancestor of Noctilionidae+Furipteridae and within Phyllostominae. The noctilionid *Noctilio leporinus* is the only piscivorous species in the superfamily, a larger-than-average species with a specialised dilambdodont dentition (Brooke, 1994; López-Aguirre, Czaplewski, et al., 2021; López-Aguirre et al., 2022). Phyllostominae includes all three specialised noctilionoid carnivores (i.e. *Chrotopterus auritus*, *Trachops cirrhosus* and *Vampyrum spectrum*), all species being of larger-than-average body size (Giannini et al., 2020; Moyers Arévalo et al., 2018). Increased molar area disparity in large carnivores reinforces the importance of increased molar and body size for the specialisation to predate on vertebrates (López-Aguirre et al., 2022).

Evolutionary shifts in dental traits were detected only within the family Phyllostomidae, signalling clade-specific processes (Supplementary Fig. 1). Decreasing surface complexity in the phyllostomid vampire subfamily Desmodontinae reflects the dental simplification in this group with adoption of a liquid diet (López-Aguirre, Czaplewski, et al., 2021; López-Aguirre et al., 2022). A shift of increasing surface complexity was detected in Phyllostominae, the only phyllostomid subfamily in which carnivory occurs (Santana & Cheung, 2016). Higher dental topographic complexity is a common adaptation in carnivorous mammals (Evans et al., 2007; López-Aguirre, Czaplewski, et al., 2021; López-Aguirre et al., 2022; Pérez-Ramos et al., 2020). Interestingly, a shift of increasing complexity was also found within the specialist hard-fruit eating subfamily Stenodermatinae, signalling increased directional selection associated with this dietary specialisation. Shifts of decreasing crown height and increasing slope were found in both Stenodermatinae and Glossophaginae, consistent with our general finding that slope and crown height had similar evolutionary trajectories. However, shifts in crown height were found at the root of each subfamily, whereas shifts in slope were found in more recent nodes within each subfamily, signalling independent patterns. The only significant shift in molar area was found in the common ancestor of the three noctilionoid carnivores *C. auritus*, *T. cirrhosus* and *V. spectrum*.

Data-driven evolutionary modelling highlighted the importance of adaptations to dietary specialisations at a finer scale than our guild categorisation. Regime shifts in multiple groups signal independent processes of directional selection in relation to adaptation to specialised diets like nectarivory, sanguivory and piscivory. Our results underscore the importance of studying the multi-faceted effect of diet on morphology at different phylogenetic scales, as well as at different ecological resolutions.

### Modes of evolution

Support for different evolutionary models across DTA metrics suggests dissociated modes of dental trait evolution in Noctilionoidea. Also, the fact that all best-supported models were diet-based, underscores the important role that diet played during noctilionoid evolution. Our OU diet-based model was best-supported for surface complexity, crown height and slope. Diet-informed BM models were the best-supported for crown sharpness and molar area, indicating no adaptive selection drove the evolution of this dental traits. Support for the BMM_diet_ model for crown sharpness and the BM_diet_ model for molar area indicates differences in ancestral states and evolutionary rates across dietary groups, but no directional selection. These results are consistent with our DTT analyses that also showed no deviation from a BM process in temporal trajectories of sharpness and molar area variation. Diet-based evolutionary models have been found to better explain the phenotypic evolution of a variety of mammal groups, including carnivorans (Law, 2021), bats (Arbour et al., 2019; Rossoni et al., 2019), primates (Aristide et al., 2018) and cetaceans (McCurry et al., 2021). Overall, our results indicate all dental traits evolved tightly linked to dietary adaptations, most traits accounting for dental shape evolving under stabilising selection (Brocklehurst & Benson, 2021), and molar area evolving under no directional selection, possibly as a result of the selective pressures that shaped body size evolution (Grossnickle, 2020).

Multivariate evolutionary modelling best supported the OU_diet_ model, indicating an evolutionary trajectory of stabilising selection towards different optima for each dietary guild. Differences between our multivariate and univariate models emphasise the importance of analysing individual traits so as not to underestimate complex evolutionary processes. The variance-covariance matrix derived for the multivariate model revealed disparate magnitudes of covariation in evolutionary rates across traits, signalling strong coevolution between crown height and curvature and surface complexity and sharpness, while also weak covariation between these two pairs of traits. Weak covariation was also found between DTA metrics of dental shape and molar area, providing more evidence for decoupled evolution of dental size and shape in Noctilionoidea. Previous studies have provided mixed evidence for patterns of covariation across DTA metrics (Berthaume et al., 2020). Although the majority of studies have focused on primates (Berthaume et al., 2019b, 2019a; Pampush et al., 2018; Ungar et al., 2017), no consensus has been reached on an overarching covariation pattern. However, it is possible that some of this covariation is partly driven by similarities in the mathematical construction of some of these DTA metrics (Berthaume et al., 2020; Thiery et al., 2019). The strong covariation between surface complexity and sharpness, and crown height and slope in this study is similar to patterns of covariation associated with changes in diet in primates (Guy et al., 2013; Ungar et al., 2017, 2018) and bats (López-Aguirre et al., 2022). Further research is needed to unveil macroecological patterns of dental morphological integration in response to diet across mammals more generally. In particular, it would be relevant to test whether the pattern of dental trait covariation found in this study is also found across other mammalian ecological and taxonomic groups.

## Conclusions

Overall, our results support our expectations of independent evolutionary trajectories across dental traits in Noctilionoidea following a mosaic evolution model (Brocklehurst & Benson, 2021). We found a marked decoupling between the evolutionary trajectories of dental shape and size, and varying degrees of evolutionary covariation between dental shape traits. Bursts of increased dental variation accompanied the heightened diversification of different phenotypic traits at different points through time. We found differences across traits in the ancestors from which modern trait diversity originated and where shifts in adaptive regimes occurred. Diversity decomposition analysis and patterns of shifts in adaptive regimes indicate that Phyllostomidae experienced the greatest dental phenotypic diversification. Differences in modes of evolution reveal dental traits evolved under different selective pressures linked to dietary specialisations. Evolutionary covariation between dental shape traits and dental size was weak, whereas covariation patterns between traits of dental shape varied. Evolutionary model testing supported different modes of evolution across dental traits, suggesting a combination of diet-based stabilising and random processes have been the main modes of dental evolution in Noctilionoidea. Moreover, the support for clade-specific processes seems to reflect the repeated evolution of dietary adaptations across Noctilionoidea, such as adaptations for frugivory in Stenodermatinae and Glossophaginae. We hypothesise that dissimilar evolutionary trajectories across dental traits enabled multiple evolutionary processes to occur in parallel, and that low trait covariation of these attributes powered the adaptive radiation of Noctilionoidea.

## Availability of data

All raw data used for this study will be uploaded as supplementary material upon acceptance for publication.

## Authors’ contributions

CL-A, MTS and SJH conceived the ideas; CL-A, GAB and MTS designed the methodology; CL-A, GAB collected and analysed the data; CL-A led the writing of the manuscript. All authors contributed critically to the drafts and gave final approval for publication.

## Acknowledgements

We thank David Boerma for his help CT scanning some of the material. GAB was funded by the BBSRC (grant BB/T01282X/1 awarded to Mario dos Reis), and CL-A is funded by a UTSC postdoctoral fellowship.

## Figure captions

**Supplementary Figure 1.** Evolutionary shifts in dental traits across Noctilionoidea. Sharpness is quantified with DNE, surface complexity with OPCR, crown height with RFI, crown curvature with slope and molar area with MA.

**Supplementary Figure 2.** Covariation in evolutionary rates across dental traits, based on the best-fitted multivariate OU_diet_ evolutionary model.

## Table Captions

**Supplementary Table 1.** Specimens list and raw data used for this study.

**Supplementary Table 2.** T tests and ANOVA tests for differences in dental topographic metrics across dietary groups. Sharpness is quantified with DNE, surface complexity with OPCR, crown height with RFI, crown curvature with slope and molar area with MA.

