## Supplementary figures and images for "Multifaceted evolution of dental morphology during the diversification of the bat superfamily Noctilionoidea"

### Supplementary Fig. 1

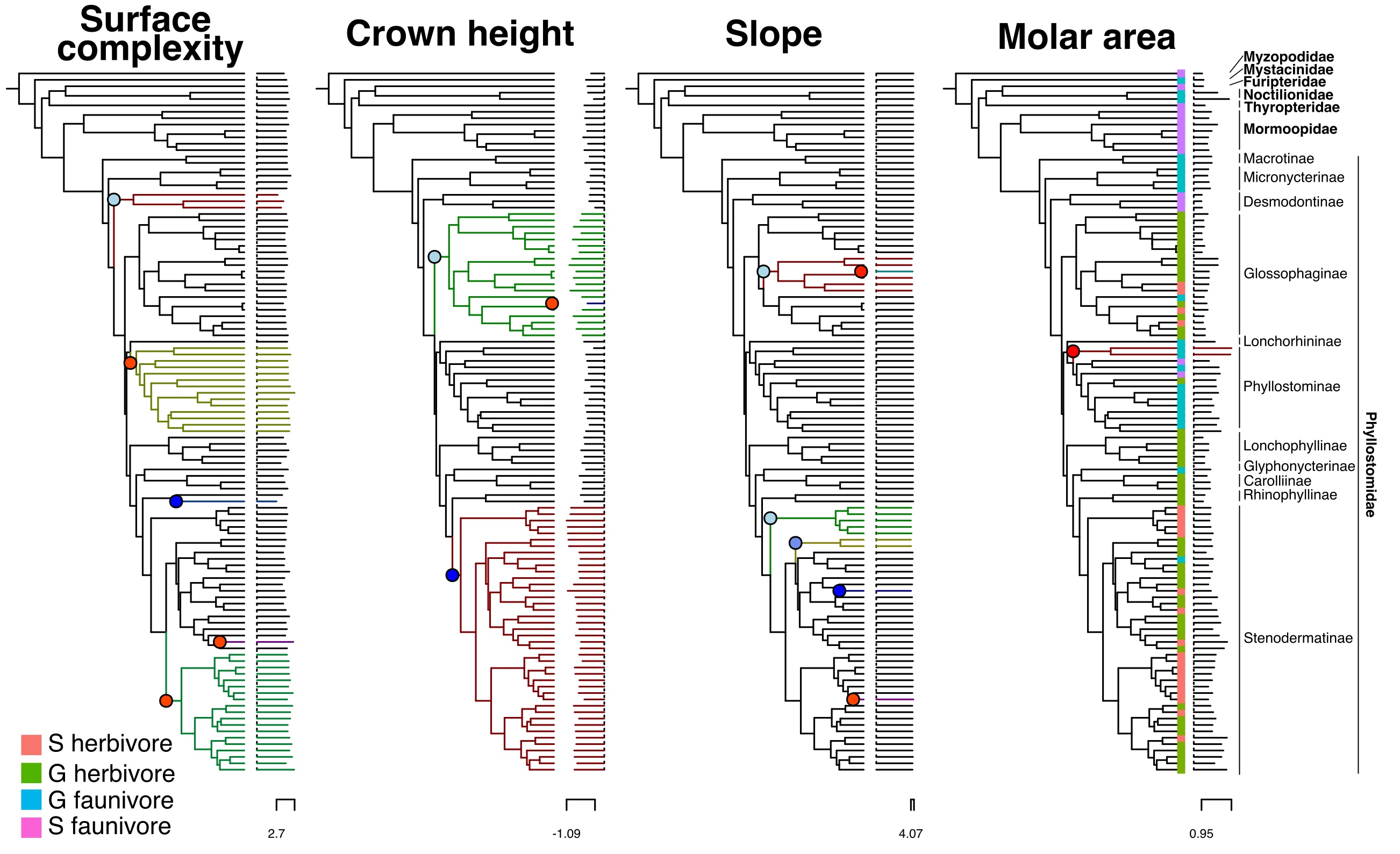

### Supplementary Fig. 2

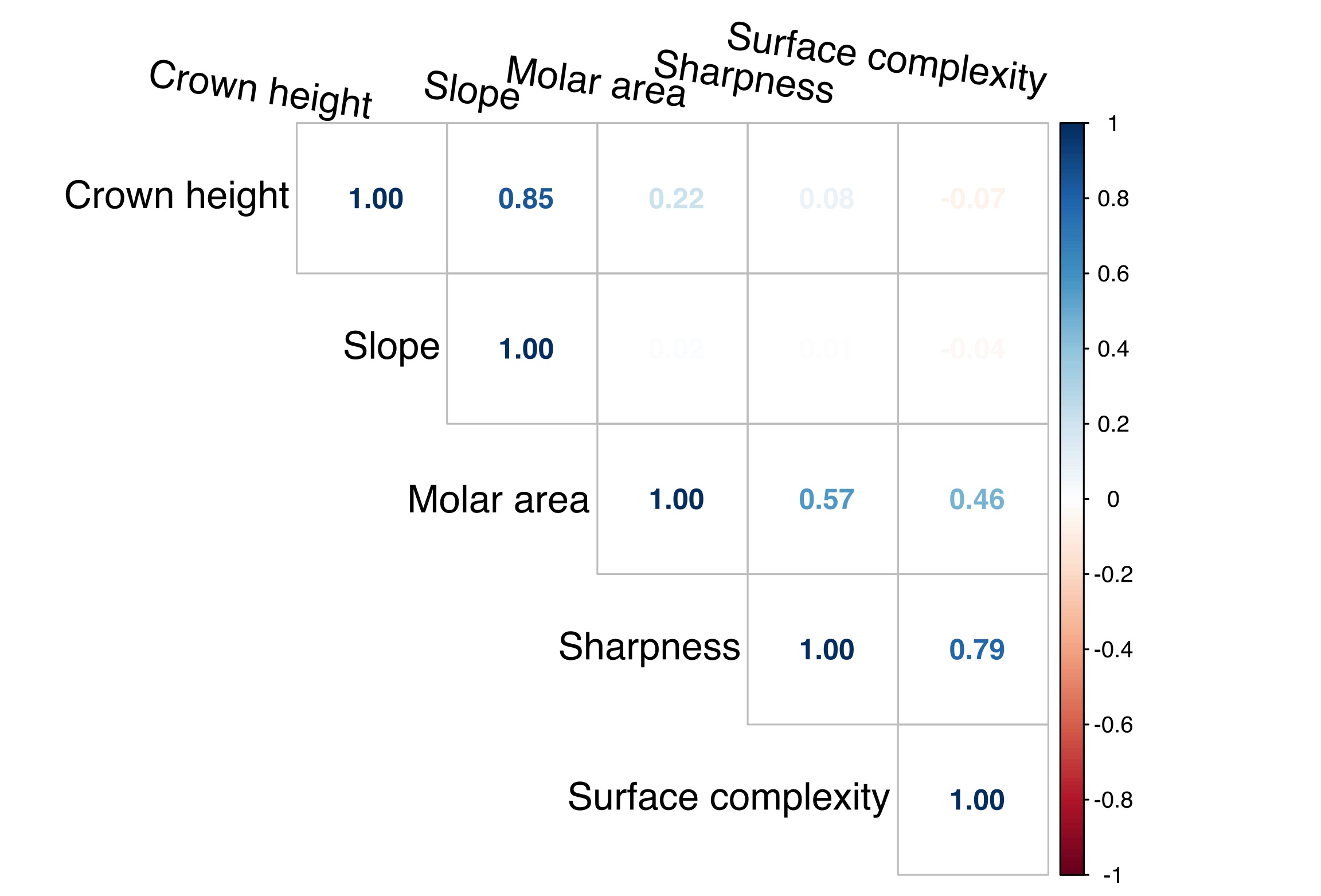
